# Trim47 inhibits murine norovirus replication in a strain-dependent manner

**DOI:** 10.1101/2025.08.08.669327

**Authors:** Stacey L. Crockett, Linley R. Pierce, Rachel Rodgers, Meagan Sullender, Lawrence A. Schriefer, Mridula Annaswamy Srinivas, Sanghyun Lee, Megan T. Baldrdige, Robert C. Orchard

**Author notes:** **Correspondence to:** **(R.C.O.)**.

## Abstract

Human norovirus is the leading cause of gastroenteritis worldwide. Norovirus exhibits remarkable genetic diversity. Understanding the impact of genetic diversity on infection and immunity has been challenging due to the difficulties of in vitro cultivation and the current lack of a small animal model. Murine norovirus (MNV) has emerged as a premier model system to investigate norovirus biology. Here, we identify Trim47 as a host restriction factor that potently inhibits MNV infection in a strain dependent manner. We determine that Trim47 expression inhibits an early stage of the viral life cycle for the MNV strain CR6, while the replication of the closely related strain CW3 is not restricted by Trim47. Using a forward genetic screen we determine that genetic variation within the nonstructural gene NS1 accounts for this differential sensitivity to Trim47. While most TRIM containing proteins promote the ubiquitination and degradation of its targets, Trim47 does neither. Instead, Trim47 promotes the deubiquitination of the NS1/2 precursor protein. Our data provide new insight into a potential antiviral gene and mechanistic insight into norovirus evolution that may impact viral tropism.

**Importance:** Viruses exist as genetically heterogeneous populations. Understanding the contribution of viral genetic variation on infection outcomes is critical in predicting emerging viruses and their variants. Noroviruses are genetically diverse but human norovirus has been technically challenging to study. In this study we use the model system murine norovirus to identify a viral strain specific restriction mechanism where a host gene can specifically restrict one strain of the virus but has no impact on a closely related strain. Dissecting the mechanism of this specificity provides insight into viral diversity and possible host restriction pathways.

## Introduction

Noroviruses are nonenveloped single-stranded RNA viruses and are a leading cause of gastroenteritis [1]. Noroviruses are genetically diverse with ten distinct genogroups, three of which infect humans [2]. Despite the robust ability of human norovirus infections to spread between humans, it has been extremely challenging to propagate human norovirus *in vitro*. Recent efforts have greatly expanded our ability to culture human norovirus, but viral replication remains limited and the ability to interrogate genetic differences between viral isolates is difficult due to a lack of a reverse genetic system for human norovirus [3].

Murine norovirus (MNV) replicates robustly in cell lines, primary cells, and in laboratory strains of mice [4]. In addition to modeling human norovirus, MNV has served as a model system for understanding viral persistence, sterilizing innate immunity, bacterial and viral interactions, development of the immune system, and microbial triggers of inflammatory diseases [5–12]. MNV is an imperfect human norovirus model as mice lack the capacity to vomit, infection rarely leads to diarrhea, and MNV encodes an additional open reading frame [4]. However, MNV and human norovirus share many significant features including genomic organization, molecular mechanisms of RNA expression and transcription, fecal-oral transmission, intestinal replication, and fecal shedding [4]. Focusing on these shared properties increases the value of studies in the MNV model as evidenced by many seminal advances in our understanding of norovirus biology using the MNV model [5–7, 12–19].

While MNV lacks the degree of genetic diversity observed in human norovirus, there is large variation in MNV infection outcomes *in vivo* [20]. For example, the prototypical MNV strains MNV^CW3^ and MNV^CR6^ share 86% nucleotide identity yet have distinct phenotypes *in vivo* [7–9, 19, 21, 22]. After oral delivery, MNV^CW3^ causes an acute infection that spreads beyond intestinal tissues and is lethal to immunodeficient mice [20]. MNV^CW3^ infects a variety of immune cells [23, 24]. In contrast, MNV^CR6^ fails to spread to systemic tissues, but infects intestinal tuft cells [19, 21]. MNV^CR6^ establishes a persistent, life-long infection in wild-type mice [7, 19, 21]. Viral chimeras between the major capsid protein (VP1) and nonstructural protein 1 (NS1) have been able to account for systemic spread and persistence, respectively [7]. However, the molecular basis for these differences in tropism remains unclear. Despite the large differences in MNV cellular tropism *in vivo*, most MNV strain differences are not observed *in vitro*. For example, both MNV^CW3^ and MNV^CR6^ can infect tuft cells *in vitro*, although only MNV^CR6^ infects tuft cells *in vivo* [25]. One exception is that MNV^CR6^ does not robustly replicate in B-cells either *in vitro* or *in vivo* [26]. Overall, our understanding of MNV strain differences has largely been limited to *in vivo* studies, creating a challenge to define the molecular mechanisms that distinguish MNV strains.

As CD300lf is a universal MNV receptor required both *in vitro* and *in vivo*, we hypothesized that unknown MNV restriction factors contribute to MNV strain specific differences [27]. We mined a previous genome-wide CRISPR activation (CRISRPa) screen identifying anti-norovirus restriction factors for MNV^CW3^ and MNV^CR6^ infection of HeLa-CD300lf expressing cells [28]. Previous attempts to identify strain specific restriction factors were unsuccessful, but one untested gene drew our attention: Trim47. Trim47 is an E3-ubiquitin ligase that is under studied and scored very strongly as an antiviral gene towards MNV^CR6^, but not MNV^CW3^ [28]. Here, we demonstrate that overexpression of Trim47 specifically restricts MNV^CR6^ but not MNV^CW3^ *in vitro*. We identify that the genetic variation within NS1 accounts for the strain specificity of Trim47. Surprisingly, Trim47 recognizes NS2 and binds the NS1/2 precursor of both MNV^CW3^ and MNV^CR6^. Trim47 expression does not lead to the degradation of NS1/2, but rather promotes the deubiquitination of NS1/2. Thus, our data point to a complicated model in which overexpression of Trim47 can restrict specific MNV strains in an NS1/2 dependent manner.

## Results

### Trim47 restricts murine norovirus replication in a strain-specific manner

Our previous data indicated that Trim47 inhibits MNV^CR6^, but not MNV^CW3^ [28]. We validated these findings by assessing the ability of MNV to replicate in HeLa cells stably expressing both CD300lf (HeLa-CD300lf) and Trim47. While MNV^CR6^ replicated robustly over 72 hours in vector control cells, MNV^CR6^ was unable to replicate above input levels in Trim47 expressing cells (**Figure 1A**). In contrast, MNV^CW3^ replication is unaffected by Trim47 expression (**Figure 1A**). Mutation of critical cysteine residues in the RING domain of Trim47 (herein called Trim47^MUT^) abrogated antiviral activity against MNV^CR6^, indicating that ubiquitin E3 ligase activity is necessary for antiviral activity (**Figure 1B**).

**Figure 1:**
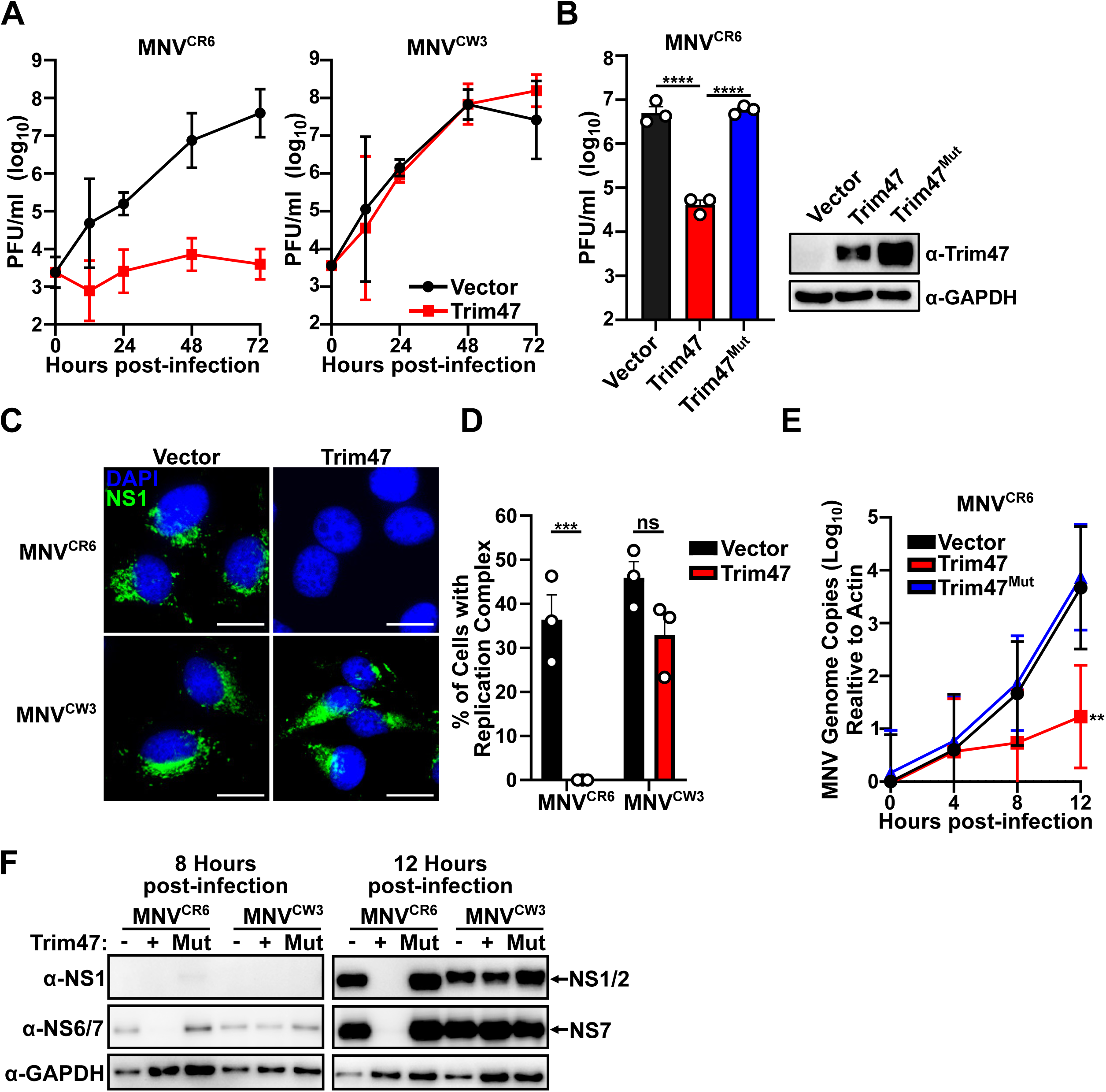
Trim47 inhibits an early step in the MNV^CR6^ but not MNV^CW3^ life cycle. **A.** HeLa-CD300lf cells expressing either an empty vector or Trim47 were infected with MNV^CR6^ (left) or MNV^CW3^ (right) at a multiplicity of infection (MOI) of 0.05. Viral production was enumerated using plaque assays at the indicated time points. **B.** HeLa-CD300lf cells expressing either an empty vector, Trim47, or Trim47^Mut^ (C9A C12A) were infected with MNV^CR6^ at an MOI of 0.05. 24 hours post-infection viral production was quantified via plaque assay. A representative western blot of indicated whole cell lysates is displayed to the right. **C-D. C)** Representative fluorescent micrographs depicting HeLa-CD300lf cells expressing either an empty vector or Trim47 infected with indicated MNV strains at an MOI of 5. Cells were stained for replication complex (NS1; green) and DAPI (blue). **D)** quantification of the percentage of cells displaying replication complexes across three independent experiments. **E.** Quantification of MNV genomes from HeLa-CD300lf cells expressing either an empty vector, Trim47, or Trim47^Mut^ infected with MNV^CR6^ at an MOI of 0.05. **F.** Representative western blots from MNV^CR6^ or MNV^CW3^ (MOI 5) HeLa-CD300lf cells expressing indicated constructs. All data are shown as mean ± S.D. from three independent experiments and analyzed by one-way ANOVA with Tukey’s multiple comparison test. Statistical significance as follows: ns not significant, \**P<*0.05, **P<0.01, \*\*\**P*<0.001, \*\*\*\**P*<0.0001

We next wanted to determine at which stage of the MNV life cycle Trim47 inhibits. We first asked if MNV^CR6^ could establish a replication complex in Trim47 expressing cells. In vector control cells both MNV^CR6^ and MNV^CW3^ formed structures positive for nonstructural protein 1 (NS1), a known marker of the replication complex (**Figure 1C**). However, in Trim47 infected cells, MNV^CR6^ was unable to form a replication complex, while formation of the MNV^CW3^ replication complex was not impeded (**Figure 1C and 1D**). Next, we assessed the ability of MNV^CR6^ and MNV^CW3^ to synthesize RNA and produce viral proteins. There was a significant reduction in MNV^CR6^ genomic copies in Trim47 expressing cells compared to empty vector and Trim47^MUT^ controls (**Figure 1E**). Similarly, while nonstructural proteins were produced in both control and Trim47 expressing cells infected with MNV^CW3^, MNV^CR6^ infection had undetectable levels of viral nonstructural proteins in Trim47 expressing cells (**Figure 1F**). These data are consistent with Trim47 inhibiting an early stage in the MNV^CR6^, but not MNV^CW3^, life cycle.

### Sensitivity to Trim47 is linked to genetic variation in NS1

To determine how genetic variation in MNV leads to different sensitivity to Trim47 restriction, we performed a directed evolution screen. We passaged MNV^CR6^ onto HeLa-CD300lf expressing Trim47 cells (**Figure 2A**). In two independent experiments MNV^CR6^ populations became resistant to Trim47 restriction after four passages (**Figure 2B**). Using our recently developed method and computational pipeline, we deep sequenced the viral population, achieving robust coverage of the viral genome (**Figure 2C**) [29]. In both experiments, a clustering of variants was found early in the viral genome corresponding to nonstructural protein NS1/2 (**Figure 2D and Supplemental Table 1**). More specifically, two mutations caught our attention due to their abundance in individual experiments: K91R and K119E (**Figure 2D**). While position 119 varies between MNV^CR6^ (Lys) and MNV^CW3^ (Arg), the identified escape mutant is a more dramatic substitution, a glutamic acid in lieu of a lysine. Additionally, we identified an association with Trim47 resistance with a substitution of lysine at position 91 for an arginine, even though both MNV^CW3^ and MNV^CR6^ encode lysine, yet differ in their sensitivity to Trim47 restriction (**Figure 2D**). To directly test if K91R or K119E confer Trim47 resistance, we introduced these mutations into our molecular clone of MNV^CR6^ and generated infectious viruses harboring individual substitutions. Both K91R and K119E mutations are sufficient to rescue MNV^CR6^ growth in Trim47 expressing cells to similar levels as control cells (**Figure 2E**). Taken together, these data point to a critical role of amino acid variants in NS1/2 in dictating MNV strain sensitivity to Trim47 mediated inhibition.

**Figure 2:**
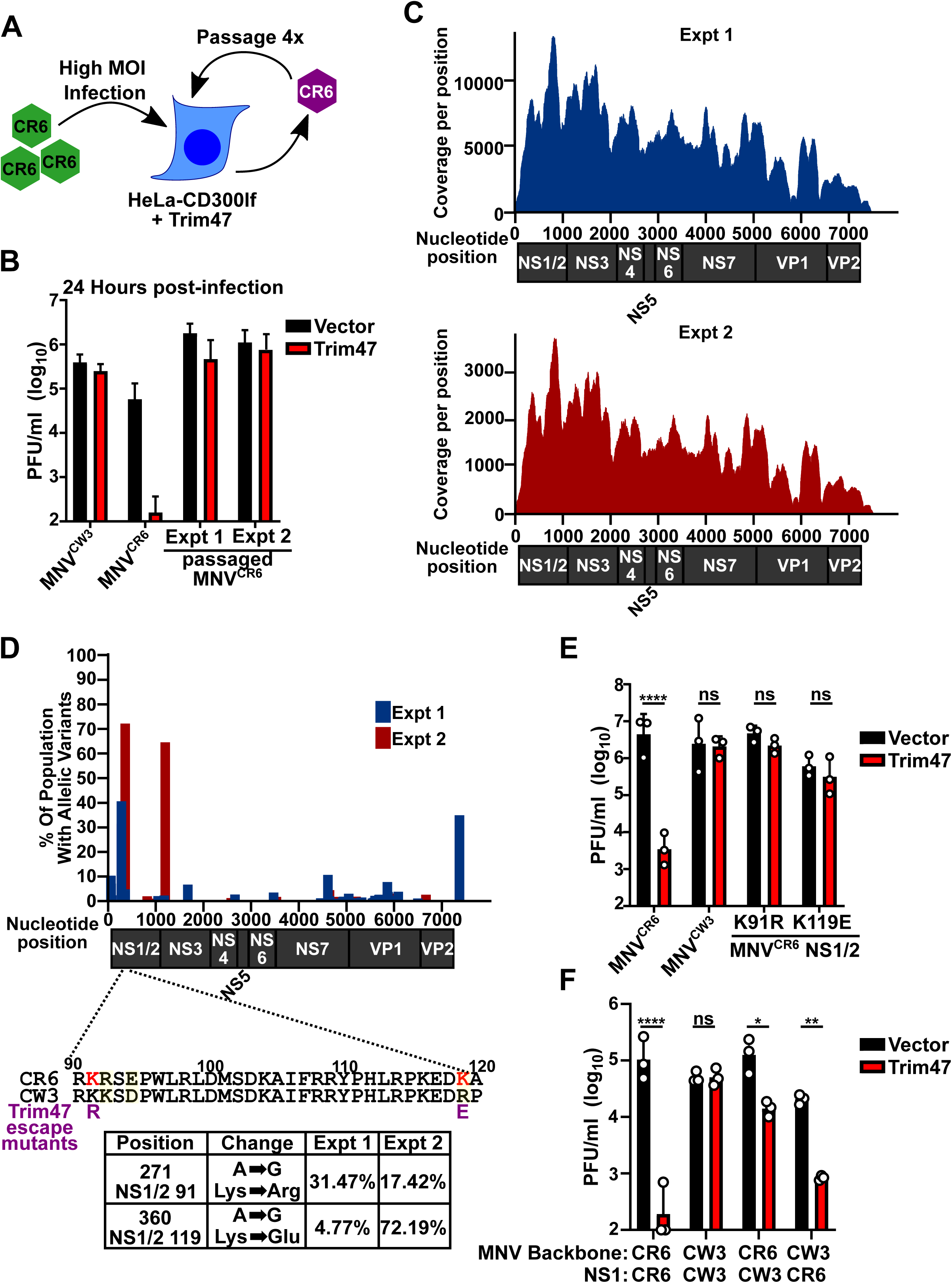
Forward genetic screen identifies variation within NS1 contributes to strain specific sensitivity to Trim47. **A.** Cartoon schematic of strategy to identify Trim47 resistance mutations. HeLa-CD300lf cells expressing Trim47 were challenged with MNV^CR6^ (MOI = 5). After 48 hours, virus was passaged onto fresh cells. **B.** HeLa-CD300lf cells expressing either an empty vector or Trim47 were infected (MOI = 0.05) with MNV^CW3^, MNV^CR6^, or MNV^CR6^ passaged onto Trim47 cells from two independent passaging experiments. Viral titers were enumerated 24 hours post-infection via plaque assay. Data are from two independent infection experiments. **C.** Summary of sequencing coverage depicting read coverage per position (y-axis) across the MNV genome (x-axis) for two independently passaged viral populations (Expt 1 and Expt 2). **D.** Summary of sequencing coverage from the two independent experiments. Graph depicts the variation as a percentage of the population (y-axis) across the MNV genome (nucleotide position; x-axis). Cartoon representation of the protein products are depicted below the nucleotide position. Below the graph is an alignment between the region of NS1 of CR6 and CW3 with differences noted by highlighting. Below the alignment are mutations enriched in the viral populations. Mutations are shown in purple below the alignment and data from the two experiments for these mutations are shown in the table. **E.** HeLa-CD300lf cells expressing either an empty vector or Trim47 were infected with indicated viruses derived from molecular clones (MOI = 0.05). 24-hours post-infection viral titers were measured via plaque assay. **F.** HeLa-CD300lf cells expressing either an empty vector or Trim47 were infected with indicated chimeric viruses derived from molecular clones (MOI = 0.05). MNV backbone refers to the origin of the virus, while NS1 indicates the specific NS1 allele in the virus. Viral titers were measured via plaque assay 24 hours post-infection. Unless otherwise indicated, all data are shown as mean ± S.D. from three independent experiments and analyzed by one-way ANOVA with Tukey’s multiple comparison test. Statistical significance as follows: ns not significant, \**P<*0.05, **P<0.01, \*\*\**P*<0.001, \*\*\*\**P*<0.0001

### Trim47 interacts with NS1/2 and NS2 in a strain-independent manner

While most MNV nonstructural proteins are processed by the viral protease, NS1/2 is cleaved by caspase 3 to generate NS1 and NS2 [30]. We next sought to understand the interactions between Trim47 and NS1, NS2, or NS1/2. For unknown reasons, exchanging the NS2 protein between MNV strains hinders the ability to recover infectious virus from molecular clones [7]. Therefore, we generated viral chimeras with the NS1 region of MNV^CR6^ and MNV^CW3^. A virus derived from MNV^CW3^ but containing the NS1 gene from the CR6 strain of MNV is sensitive to Trim47 inhibition indicating that CR6 NS1 is sufficient to convert a resistant virus to being sensitive to Trim47 (**Figure 2F**). Introduction of the NS1 sequence from the CW3 strain of MNV into the backbone of MNV^CR6^ partially rescued viral replication (**Figure 2F**). These data highlight that the genetic variation within the NS1 region is a critical determinant of Trim47 sensitivity. However, due to the incomplete rescue of replication for MNV^CR6^ containing the NS1 from MNV^CW3^, these data also point to a role for other genomic features that may contribute to MNV strain selectivity of Trim47.

Given the importance of NS1 in determining MNV sensitivity to Trim47-mediated restriction, we tested whether Trim47 interacts with NS1. While Trim47 co-immunoprecipitated with NS1/2, we did not detect any interactions with NS1 (**Figure 3A**). Surprisingly, we detected a robust co-immunoprecipitation between NS2 and Trim47 (**Figure 3A**). We also detected a physical interaction between Trim47 and the NS1/2 protein from both MNV^CR6^ and MNV^CW3^ (**Figure 3B**). Furthermore, MNV^CR6^ NS1 protein containing the escape mutants K91R and K119E still co-immunoprecipitates with Trim47 (**Figure 3B**). These biochemical interactions point to a role beyond physical binding in determining the sensitivity of MNV strains to Trim47 antiviral activity.

**Figure 3:**
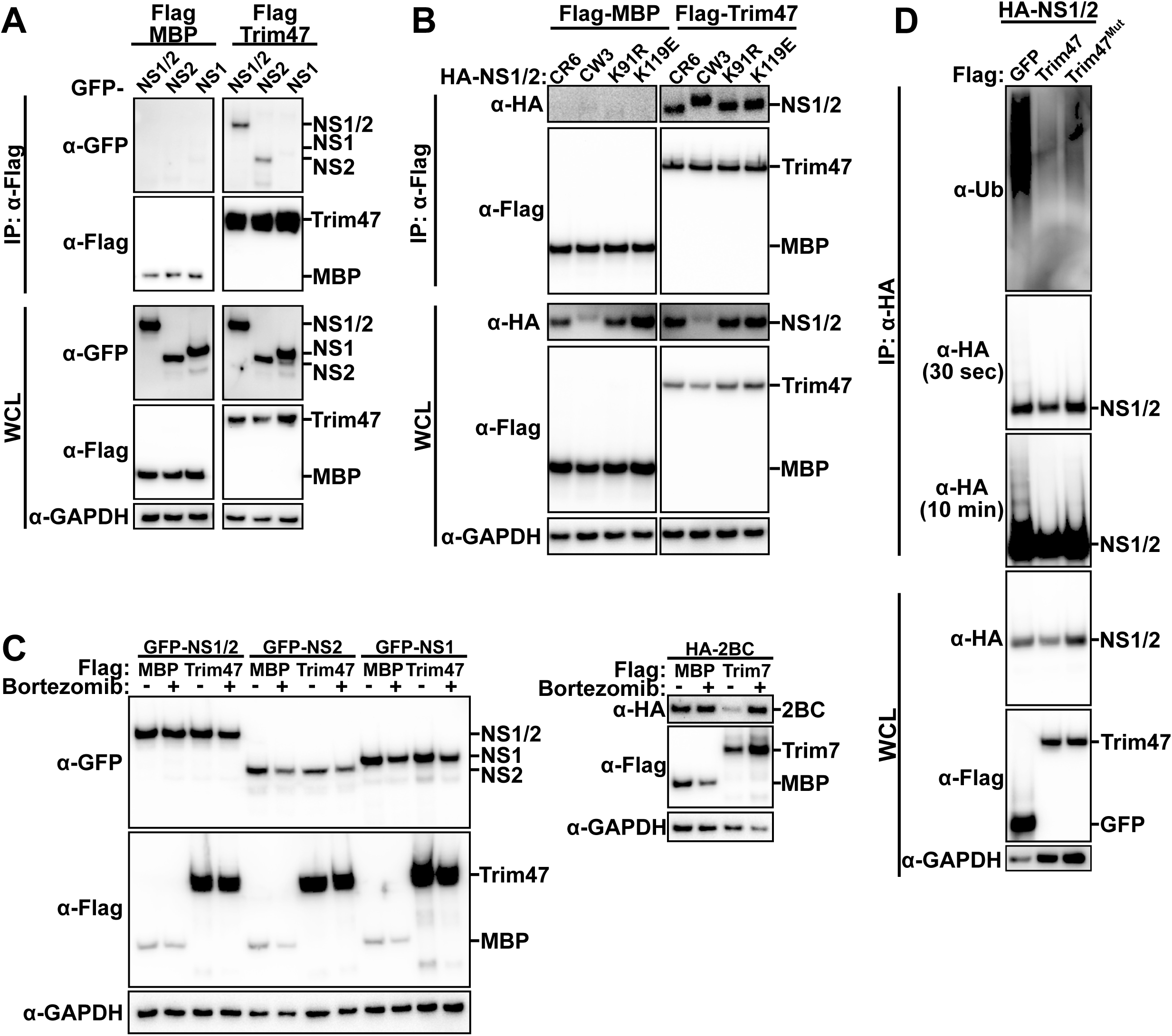
Reconstitution of the Trim47 and NS1/2 interactions. **A.** Representative western blot from co-immunoprecipitation experiment in which HEK293T cells were co-transfected with GFP-tagged NS1/2, NS2, or NS1 along with Flag-tagged MBP or Trim47. Whole cell lysates (WCL) were subjected to anti-Flag immunoprecipitation. **B.** Co-immunoprecipitation analysis from transfected HEK293T cells of indicated HA-tagged NS1/2 constructs with Trim47. **C.** Western blot analysis of HEK293T cells co-expressing indicated proteins. 24 hours post-transfection, cells were treated with either DMSO or 100nM Bortezomib (Sigma 5043140001) for an additional 8 hours. Data on the right with Trim7 and Coxsackievirus 2BC was conducted concurrently with the Trim47 and NS1/2 degradation experiment and is provided as a positive control for the assay **D.** In cell ubiquitination assay using transfected HEK293T cells with indicated constructs and immunoprecipitated for HA-NS1/2. All data are representative images from three independent experiments.

Many antiviral TRIM domain containing proteins function via degrading viral proteins [31]. However, co-expression of Trim47 did not promote proteasome-mediated degradation of NS1, NS2, or NS1/2 (**Figure 3C**). Importantly, as an experimental control we detected proteasomal degradation of the Coxsackievirus B3 protein 2BC by Trim7 in the same experiment (**Figure 3C**) [32]. While degradation is a common fate for proteins tagged with ubiquitin, there are degradation independent functions of antiviral TRIM containing proteins [31]. We set out to determine if Trim47 ubiquitinates the NS1/2 of MNV^CR6^ when co-expressed in 293T cells. We found substantial ubiquitination of NS1/2 when co-expressed with the control protein GFP (**Figure 3D**). Contrary to our hypothesis, co-expression with Trim47 decreased NS1/2 ubiquitination (**Figure 3D**). This reduction in ubiquitination is largely, but not completely, dependent upon a functional RING domain as expression of Trim47^MUT^ only modestly reduces NS1/2 ubiquitination levels. Unfortunately, we were unable to perform the degradation and ubiquitination experiments during infection as MNV^CR6^ viral protein levels are undetectable in Trim47 expressing cells (**Figure 1F).** Also, we confirmed that the addition of proteasome inhibitors blocks the replication of MNV independently of Trim47 [33]. Thus, it is possible that Trim47 mediated ubiquitination or degradation of MNV non-structural proteins only occurs in the context of MNV infection. However, these data suggest that the antiviral mechanism of Trim47 may be due to the decrease in ubiquitination of NS1/2. Taken together, these data point to a complicated, noncanonical mechanism by which Trim47 specifically restricts MNV^CR6^ but not MNV^CW3^ infection.

### Processing of NS1/2 does not explain MNV strain specificity of Trim47

We next tested a model in which MNV strains or escape mutants differ in their sensitivity to Trim47 due to different abilities to process NS1/2 into NS1 and NS2. First, we tested the hypothesis that caspase 3 processing of NS1/2 promotes Trim47 sensitivity. MNV^CR6^ virus harboring a pair of mutations (D121G and D131G) that do not hinder viral propagation but block caspase mediated cleavage of NS1/2 (herein called MNV^CR6^ΔCasp) was equally inhibited by Trim47 as the parental virus (**Figure 4A)** [17, 30, 34]. Next we explored the opposite hypothesis where enhanced processing of NS1/2 by caspase 3 increases resistance to Trim47 restriction. In support of this hypothesis, the K119E escape mutant falls within the first caspase 3 cleavage site and is predicted to increase cleavage [35]. Indeed, in the absence of Trim47 MNV^CR6^ harboring the Trim47 escape mutant K119E has an increase in NS1/2 processing compared to parental MNV^CR6^ or the other Trim47 escape mutant MNV^CR6^ NS1-K91R mutant (**Figure 4B**). To directly test if an increase in NS1/2 processing contributes to the resistance of MNV strains to Trim47, we added the pan-caspase inhibitor zVAD during infection of control or Trim47 overexpressing cells. Addition of zVAD blocked the processing of NS1/2 of all MNV viruses tested similarly to the MNV^CR6^ΔCasp (**Figure 4C**). Despite this block in NS1/2 processing, MNV^CW3^ and both Trim47 escape mutants (MNV^CR6^ harboring either K91R or K119E) remained resistant to Trim47 inhibition in the presence of zVAD (**Figure 4D**). Taken together, our data argues against a major role for NS1/2 processing by host caspases in determining sensitivity to Trim47.

**Figure 4:**
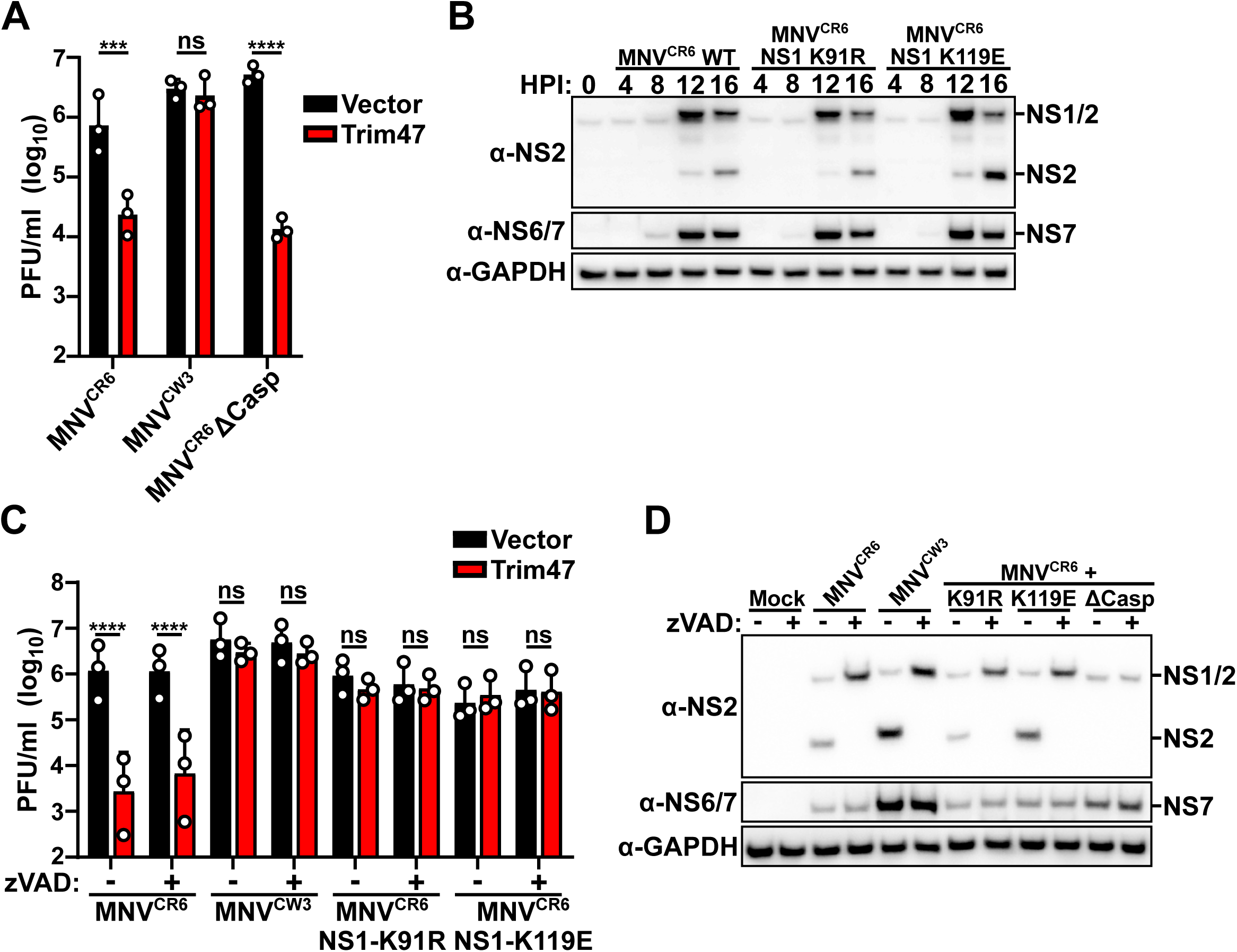
Caspase processing of NS1/2 does not contribute to different Trim47 sensitivity of MNV strains. **A.** HeLa-CD300lf cells expressing either an empty vector or Trim47 were infected with MNV^CW3^, MNV^CR6^, or MNV^CR6^ΔCasp (NS1 D121G, D131G) at an MOI of 0.05. 24 hours post-infection viral titers were enumerated via plaque assay. **B.** A representative western blot from BV2 cells infected with indicated MNV strains at an MOI of 5 for different hours post-infection (HPI). Expected molecular weights of NS1/2 and NS2 are indicated. **C-D)** HeLa-CD300lf cells expressing either an empty vector or Trim47 were infected with indicated MNV strains in the presence of vehicle (-) or zVAD (+). 24 hours post-infection viral titers were enumerated **(C)** or polyprotein processing assessed via western blot **(D).** All data are shown as mean ± S.D. from three independent experiments and analyzed by one-way ANOVA with Tukey’s multiple comparison test. Statistical significance as follows: ns not significant, \**P<*0.05, **P<0.01, \*\*\**P*<0.001, \*\*\*\**P*<0.0001

## Discussion

Despite significant variation in cellular and tissue tropism of MNV strains during *in vivo* infections, a molecular understanding of these differences has been missing due to the near uniform growth of MNV strains *in vitro*. Here we identify an MNV strain-specific growth phenotype in cells overexpressing Trim47. Interestingly, we mapped the genetic basis of Trim47 strain dependent inhibition of MNV to NS1 (**Figure 2)**. NS1 is a gene whose genetic variation is necessary and sufficient to explain tuft cell tropism of persistent MNV strains *in vivo* [7, 17–19, 25]. It is important to note that the NS1 variants that promote persistence also render MNV sensitive to Trim47 (**Figure 2F**). We identified a discrepancy between the genetic sensitivity to Trim47 and the biochemical interactions of Trim47 and viral proteins. Genetic variation within NS1 largely, but incompletely, accounts for Trim47 strain dependent phenotypes (**Figure 2D-2F**). Biochemical interactions between Trim47 and NS2 were robust while an interaction with NS1 and Trim47 was undetectable (**Figure 3A**). Trim47 did interact with the precursor protein NS1/2. NS1/2 is unique amongst noroviruses as it is cleaved by host caspases rather than the viral protease [30]. We did not detect a significant impact of enhancing or eliminating NS1/2 processing on Trim47 mediated restriction (**Figure 4**). The distinct mechanism of NS1/2 processing suggests some unique relationship between NS1 and NS2. Our data suggests that this relationship might be at the heart of Trim47 mediated inhibition as genetics and biochemistry implicate different regions of NS1/2. However, future work defining whether NS1 and NS2 cooperate as individual proteins or as domains in the NS1/2 precursor is necessary to enhance our understanding of norovirus biology. Furthermore this insight is likely to help explain how Trim47 selectively restricts MNV^CR6^ but not MNV^CW3^ replication.

The antiviral activity of Trim47 requires a functional E3 ligase domain (**Figure 1B**). In reconstitution studies we did not detect ubiquitination of NS1/2 by Trim47. Rather NS1/2 is ubiquitinated in the absence of Trim47 overexpression in 293T cells (**Figure 3D**). Surprisingly, Trim47 co-expression decreased ubiquitination of NS1/2. It remains unclear how Trim47 expression decreases NS1/2 ubiquitination. One possibility is that Trim47 catalyzes the addition of a ubiquitin-like molecule rather than ubiquitin. Some TRIM proteins can also function as E3 SUMO ligases [36]. For example, Trim38 can polyubiquitinate some substrates such as TRIF, but Trim38 SUMOylates MDA5, RIG-I, and cGas [37, 38]. Interestingly, the SUMOylation of MDA5 and RIG-I by Trim38 inhibits their polyubiquitination [37]. Whether Trim47 catalyzes a ubiquitin-like modification to block polyubiquitination is an intriguing possibility but needs experimental validation. Trim47 may use an alternative method to counter the ubiquitination of NS1/2 by other E3 ligases. This may occur through direct or indirect methods such as degrading the bona fide NS1/2 ubiquitin ligase or recruiting a deubiquitinase (DUB) to NS1/2. To this latter point, TRAF6, an E3 ubiquitin ligase, also recruits the DUB CYLD to substrates leading to the loss of ubiquitination on target proteins [39]. Interestingly in addition PKC-ε and NF-90, CYLD is a reported Trim47 interacting protein and substrate of Trim47 ubiquitination [40, 41]. However, there is no obvious connection between these host proteins and MNV replication. The impact of NS1/2 ubiquitination for norovirus replication or for the function of NS1/2 is not clear. In the future, determining whether NS1/2 is ubiquitinated during infection will enhance our understanding of MNV biology and the mechanism by which Trim47 inhibits viral replication.

Our study has several limitations. First, our data relies on overexpression of Trim47 and we do not assess the role of endogenous Trim47. The advantages of a near binary viral phenotype enabled us to define mechanistic differences between MNV strains but may not represent the physiological role of Trim47. Thus, future studies leveraging genetic loss of function including in mice will be necessary to define the physiological role of Trim47. Additionally, our mechanistic and biochemical studies utilized reconstitution studies since MNV^CR6^ protein levels were undetectable in Trim47 expressing cells. The lack of NS1/2 degradation or ubiquitination induced by Trim47 may be a result of a missing component in our reconstituted system. Lastly, while we identified multiple potential escape mutants within the MNV polypeptide, we only focused on two escape mutants. While each mutant was sufficient to confer resistance to viral infection, we could not detect any biochemical differences between these escape mutants and the wild-type proteins. It is likely that their mechanisms may only be revealed in the context of infection rather than our reconstitution studies. Nevertheless, the strain dependent restriction of MNV by Trim47 *in vitro* provides an opportunity to learn about norovirus genetic diversity and perhaps gain information on the function of the poorly understood viral nonstructural proteins NS1, NS2, and their precursor NS1/2.

## Materials and Methods

### Cell Culture

293T (ATCC), BV2 cells (kind gift of Dr. Skip Virgin, Washington University), and HeLa cells (ATCC) were cultured in Dulbecco’s Modified Eagle Medium (DMEM) with 5% fetal bovine serum (FBS). Stable cell lines were generated by lentivirus transduction. Briefly, lentiviral vectors were co-transfected with packaging vector (psPax2) and pseudotyping vector (pCMV-VSV-G) into 293T cells using Transit-LT1 (Mirus). 48 hours post-transfection, lentivirus was collected, filtered through a 0.45 µm filter, and added to cells. 48 hours post-transduction, media was changed to contain the appropriate antibiotic (5 µg/ml blasticidin and/or 1 µg/ml puromycin). All cell lines are tested regularly and verified to be free of mycoplasma contamination.

### Plasmids

Human Trim47 cDNA was cloned into pCDH-MSCV-T2A-Puro vector and pCMV-N-Flag (Clontech) for lentiviral and transient expression respectively. For reconstitution studies MBP and eGFP were cloned into pCMV-N-Flag. MNV^CR6^ NS1/2 (1-341), NS2 (132-341), and NS1 (1-131) were cloned downstream of eGFP in pcDNA3.1. The NS1/2 sequence of MNV^CR6^ and NS1/2 of MNV^CW3^ were cloned in pcDNA3.1 HA (Addgene #128034). Molecular clones for MNV^CW3^ (GenBank accession EF014462.1) and MNV^CR6^ (GenBank accession JQ237823) have been described previously [42]. Point mutations including NS1-K91R, NS1-K119E, and Trim47 C9AC12A were introduced via splicing by overlap extension PCR. All plasmid sequences were verified through sanger sequencing prior to use.

### Viral Assays

MNV^CW3^, MNV^CR6^, and respective mutants were generated by transfecting molecular clones into 293T cells and amplifying on BV2 cells as described previously [43]. For growth curves, 5 x 10^4^ indicated HeLa-CD300lf cells were seeded in 96-well plate and subsequently infected with MNV strains at a multiplicity of infection (MOI) of 0.05. Samples were subsequently frozen at −80°C at the indicated time points. Viral titers were enumerated via plaque assay as described previously [43].

For immunofluorescence microscopy, HeLa-CD300lf expressing either an empty vector or Trim47 were seeded overnight onto glass coverslips in six-well dishes. The next day, cells were infected with MNV^CW3^ or MNV^CR6^ at an MOI of 5. 24 hours post-infection, cells were washed with ice-cold PBS, fixed with 4% PFA, and permeabilized with 0.5% Triton X-100, blocked with 1% BSA and stained with mouse anti-NS1 and anti-mouse AlexaFluor 488 (ThermoFisher) and mounted using Prolong Gold with DAPI. For each of the three independent experiments, five randomized images containing at least 20 cells were scored for the presence or absence of a replication complex.

For quantification of viral genomes, indicated HeLa-CD300lf cells were infected with MNV^CR6^ at a MOI of 0.05 and TRI Reagent (Sigma-Aldrich #T9424) were added at the indicated time points. RNA was isolated using the Direct-zol RNA Prep kit (Zymo Research) following the manufacturer’s protocol. Purified RNA was used for cDNA synthesis using the High-Capacity cDNA Reverse Transcription kit (Thermo Fisher Scientific #4368813). TaqMan quantitative PCR (qPCR) for MNV was performed in triplicate on each sample and standard with forward primer 5’-GTGCGCAACACAGAGAAACG-3’, reverse primer 5’-CGGGCTGAGCTTCCTGC-3’, and probe 59-6FAM-TAGTGTCTCCTTTGGAGCACCTA-BHQ1-3’. TaqMan qPCR for actin was performed in triplicate on each sample and standard with forward primer 5’-GATTACTGCTCTGGCTCCTAG-3’, reverse primer 5’-GACTCATCGTACTCCTGCTTG-3’, and probe 5’-6FAM-CTGGCCTCACTGTCCACCTTCC-6TAMSp-3’.

For detecting viral protein production, cells were infected with indicated MNV strains at an MOI of 5 and lysed in RIPA buffer containing HALT protease and phosphatase inhibitors at indicated time points. Lysates were clarified via centrifugation prior to Western Blot analysis.

### Directed Viral Evolution

These forward genetic experiments with MNV were performed under BSL2 conditions, as approved by the UT Southwestern Institutional Biosafety Committee. Two independent viral passaging experiments were conducted in HeLa-CD300lf Trim47 expressing cells using a similar strategy as we have described previously [44]. Briefly, 1 x 10^6^ HeLa-CD300lf Trim47 expressing cells were seeded in a 10-cm2 plate and subsequently infected with MNV^CR6^ at an MOI of 5. 48 hours post-infection, supernatants from the cultures were harvested and clarified (10 minutes at 3,000 x *g*). 1 ml of the clarified supernatant was added onto 1 x 10^6^ HeLa-CD300lf Trim47 expressing cells were seeded in a 10-cm^2^ plate. This passaging was performed four times and 1 mL of clarified supernatant was used to isolate total RNA using the Direct-zol RNA Prep Kit (Zymo Research). Sequencing and data analysis were performed identical to what we have previously described [29, 44].

### Antibodies and Western Blotting

Samples were subjected to SDS-PAGE and subsequently transferred to PVDF membranes. Membranes were blocked in TBS-T supplemented with 5% non-fat dry milk prior to probing with antibodies. Antibodies used include: rabbit α-Trim47 (1:1000; Abcam: ab72234), mouse α-GAPDH-HRP (1:10,000; Sigma: G9295), Rabbit α-GFP (1: 2,000; Invitrogen A-11122), mouse α-FLAG M2-HRP (1:2500; Sigma: A8592), Rabbitt α-HA (1:1,000; Cell Signaling: C29F4), mouse α-Ubiquitin-HRP (1 µg/ml; Cytoskeleton: AUB01-HRP), α-Rabbit-HRP (1:10,000; ThermoFisher: 34102), and α-mouse-HRP (1:10,000; Sigma: SAB3701122). Antibodies for MNV non-structural proteins including mouse α-NS1, rabbit α-NS2, and rabbit α-NS6/7 were used as described previously [45].

### Coimmunoprecipitation

293T cells were seeded at 1 x 10^6^ cells per well of a six well plate and transfected with indicated plasmids. At 24 hours post-transfection cells were lysed in RIPA Buffer (10mM Tris [pH 7.5], 140mM NaCl, 1mM EDTA, 0.5mM EGTA, 0.1% DOC, 0.1% SDS, 1% Triton X-100) containing HALT protease and phosphatase inhibitors. Clarified lysates were incubated with anti-FLAG-M2 (1:500; Sigma F1804) with gentle rocking at 4°C. 2 hours later, Protein A/G Agarose (ThermoFisher 20421) was added to lysates and incubated overnight with gentle rocking at 4°C. Samples were subjected to four washes with RIPA buffer and proteins were eluted from beads with Laemmli buffer and subjected to SDS-PAGE and western blotting as described above with the addition of the TidyBlot Western Blot Detection Reagent (Bio-Rad STAR209PT)

### Degradation Assay

293T cells were seeded at 2 x 10^5^ per well in a 24-well plate and subsequently transfected with 250ng of the indicated plasmids. After 24 hours, cells were treated with either DMSO or 100nM Bortezomib (Sigma 5043140001) for an additional 8 hours. Cells were lysed in Laemmli buffer and subjected to SDS-PAGE and western blotting.

### Ubiquitination of NS1/2

293T cells were seeded at 1 x 10^6^ per well of a six-well plate and transfected with corresponding plasmid construct. At 24 hours post-transfection, cells were treated with 100 nM Bortezomib for an additional 8 hours. Cells were then lysed in RIPA Buffer containing HALT protease and phosphatase inhibitor cocktail and lysates were incubated with anti-HA-agarose (Sigma A2095) overnight on a nutator at 4°C. The next day, samples were subjected to four washes with RIPA buffer and proteins were eluted from beads with Laemmli buffer and subjected to SDS-PAGE and western blotting.

### Inhibition of NS1/2 processing

To verify that the caspase inhibitor z-VAD (OMe)-FMK (Cell Signaling Technology 60332S) effectively prevents NS1/2 processing, HeLa-CD300lf cells were seeded at 2.5 x 10^5^ per well in a 12-well plate and infected the next day with indicated MNV strains at an MOI of 5. 8 hours post-infection, cells were treated with either DMSO or 50μM Z-VAD. 24 hours post-infection, cells were lysed in Laemmli buffer and analyzed via SDS-PAGE and western blotting. For MNV growth assays in the presence of the caspase inhibitor z-VAD, HeLa-CD300lf cells expressing either an empty vector or Trim47 were infected with indicated MNV strains at an MOI of 0.05. 8 hours post-infection, cells were treated with either DMSO or 50μM Z-VAD. Samples were frozen at −80°C 24 hours post-infection and infectious virus was measured via plaque assay.

## Acknowledgements

We would like to thank Craig Wilen, Kim Orth, Julie Pfeiffer, John Schoggins and all members of the Orchard lab for helpful discussions on this project. S.L.C. was supported by a UT Southwestern Immunology T32 Training Grant (AI005284). R.C.O. was supported by the NIH (5R01DK133231) and by the Welch Foundation (I-2006-20190330).

## Author Contributions

S.L.C. designed the project, performed experiments, and helped to draft the paper. L.R.P. helped to design the project and perform experiments. R.R., M.E.S., L.A.S. M.S. performed experiments. S.L. provided valuable reagents and conceptualization of the project. M.T.B. helped conceptualize the project, provided supervision, and data analysis. R.C.O. conceptualized the project, provided supervision, analyzed data, and wrote the paper. All authors read and edited the manuscript.

